# 3-Mercapto-3-methyl-1-butanol, a component of leopard urine, repels leopards and spotted hyaenas and protects livestock from leopard attacks; a series of case studies

**DOI:** 10.1101/2025.09.14.676068

**Authors:** Peter Apps, Peter Makumbe, J.W. McNutt

## Abstract

Mammals’ semiochemicals could provide tools to control pests, deter unwanted behaviours, enhance the survival of imperilled species, and improve the productivity and welfare of domestic animals, but because producing facsimile copies of whole odours is unrealistic in terms of technical challenge and cost, the real-world application of mammals’ chemical signals depends on mammals responding to simple subsets of the components of their social odours.

Experiments in multiple locations in southern Africa have shown that the odour of 3-mercapto-3-methyl-1-butanol (3M3MB), a component of leopard urine, can protect livestock from leopards, and repel spotted hyaenas.

Camera-trapping for four-months at an overnight enclosure (known locally as a kraal), on a cattle ranch in Botswana, with no 3M3MB being released, yielded seven records of leopards and seven of spotted hyaenas, and one calf was killed by a leopard. Over the following 4.5 months, with 3M3MB being released, there were no records of leopards, one record of spotted hyaenas, and no losses of calves.

At another ranch, after three leopards entered through a gate during a one-month control period, 3M3MB was deployed at increasing release rates, and the number of leopard records decreased to zero as the release rate increased, but then rose again after five months of continuous use. On the same ranch, when sheep and goats were fitted with collars emitting 3M3MB, losses to leopards declined from seven per month to zero for the five months that the collars were in operation.

On a third Botswana ranch, leopards killed 0.6 calves per month when no 3M3MB was present and 0.2 calves per month when 3M3MB was released. The value of the livestock saved was approximately six times the cost of the repellent.

At Shangani, in Zimbabwe, camera traps captured images of two leopards just outside shade-cloth kraals where 3M3MB was being released, but no livestock were attacked despite the shade cloth fence providing no physical barrier. On the same ranch, a leopard killed 4 calves in 5 days at a traditional thorn-branch kraal, 3M3MB and camera traps were deployed for the following month, during which no leopard images were captured nor calves killed. After the 3M3MB was removed, 2 calves were killed within 10 days.

3M3MB deterred spotted hyaenas from crossing a veterinary cordon fence in Botswana, and its efficacy rose with release rate.

## 1. Introduction

Just as with insects, mammals’ behaviour and physiology are influenced by chemical signaling between individuals. The potential application of these chemical signals to pest control, breeding or conservation was recognized in the early 1980s (summary in Apps and McNutt 2023), but has remained unfulfilled, in large part because the entrenched orthodoxy that mammalian chemical signals are coded by irreducibly complex mixtures (Albone 1984, Pickett et al. 2014; Müller-Schwarze 2016) has been a barrier both to the identification of signaling compounds (Apps 2013), and to their practical application, because such complex mixtures would be prohibitively costly to produce and apply. To bring chemical signaling into practical use depends on mammal behaviour or physiology being manipulated by single compounds or simple mixtures of compounds.

Insect chemical signals (pheromones) have been effective in the control of insect pests, and had a market value of nearly $4 Billion in 2023, which is projected to grow to nearly $11 Billion by 2029 https://www.fortunebusinessinsights.com/industry-reports/agricultural-pheromones-market-100071 (accessed 13 Jan 2025). In stark contrast, the potential of mammal chemical signals for reducing the damage caused by mammalian crop pests and livestock-predators, and protecting endangered wildlife remains unrealized, and commercial mammal pheromones are limited to artificial boar odor to test estrus in sows (*Sus scrofa domesticus*), and “appeasing pheromones” for pets and farm animals. There are currently no commercial applications of mammal chemical signals to reduce the billions of dollars in crop losses inflicted by rodents (which are also vectors and reservoirs of zoonotic diseases) (Hohenberger et al. 2022; RoDiagne et al.2023), or to protect the billions of dollars’ worth of livestock killed by predators.

Here we report on experimental field tests of the effects of a single component of leopard (*Panthera pardus*) urine; 3-mercapto-3-methylbutanol (3M3MB) (Apps et al. 2014), on leopard predation of livestock and the movements of spotted hyaenas (*Crocuta crocuta*) on commercial livestock ranches in Botswana and Zimbabwe and subsistence cattle posts in northern Botswana. The results provide proof of the concept that a technically simple formulation of a single component of a mammal chemical signal has a practical, commercially viable application.

### 1.1 Human-predator conflict

There is a global decline in the numbers of wild predators and the viability of their populations, as their habitats are lost to expanding human populations with their livestock and crops (Fernandez-Sepulveda and Martin 2022). Conflict with humans is the single biggest threat to predator populations worldwide, but at the same time, depredation on livestock has significant negative impacts on rural livelihoods, especially on subsistence pastoralists with small flocks of sheep (*Ovis aries*) and goats (*Capra hircus*). In South Africa, predators kill livestock worth ZAR1 billion each year (Kerley et al. 2017), and in the USA losses of small stock to predators cost $US40 million, and of cattle $US98.5 million *per annum* (USDA 2010, 2011). In Shorobe, a traditional livestock village in NW Botswana, 67% of livestock owners reported annual losses of up to 14% of goats and 3% of cattle to predators (McNutt et al. 2017), and in the Kalahari in Botswana some cattle posts suffer more than 6 depredations per month (Rutina et al. 2017).

As long as it is easier and cheaper for farmers to kill predators than to use non-lethal methods such as herding, predator-proof bomas, or early-warning systems, lethal predator control will continue. Globally, habitat loss and the lethal control that accompanies it have extirpated some species of large carnivores from more than 90% of their historical range (Wolf and Ripple 2017), and small and mesocarnivores at regional scale (Marneweck et al. 2021). Indiscriminate lethal control methods such as gin-traps, snaring and poisoning often kill non-target species (McManus et al. 2015), and in Africa, non-target poisoning of vultures is a particular concern (Nyirenda et al. 2024).

Official records of lethal control underestimate its true scale because “shoot, shovel and shut up” is common. Lethal control is the first option for > 21% of cattle posts in Shorobe (McNutt et al. 2017), and a large majority of the Shorobe farmers report that they would “remove” (in other words kill) a predator in response to loss of livestock. Livestock owners’ aversion to predators is due to predators killing livestock, and where attacks on livestock are fewer, tolerance for predators is higher (Lindsey et al. 2013). Non-lethal methods of preventing predation and protecting rural livelihoods without adverse ecological effects (McManus et al. 2015), need to be economically sustainable and applicable to local circumstances, but hardly any existing non-lethal interventions meet these criteria. Early warning systems based on satellite tracking of collared predators, intensified herding and husbandry (Hawkins et al. 2023), and compensation (Littlewood et al. 2020; Montag 2023), all fall short due to costs (McManus et al. 2015; Weise et al. 2018), culturally embedded practices (McNutt et al. 2017; Mmopelwa & Mpolokeng 2008), and lack of motivation (Mmopelwa & Mpolokeng 2008). In Botswana, local breed livestock guarding dogs (*Canis familiaris*) are very effective protection for goats and sheep (Van der Weyde et al. 2020; Horgan et al. 2021) and are the only non-lethal intervention with any spontaneous uptake in the subsistence sector. In South Africa LGDs are more effective and more cost effective than lethal control (McManus et al. 2015).

Predator conflict prevention and mitigation is stuck in a rut of ineffective compensation schemes, unsustainable subsidies to livestock husbandry, and high-tech interventions that need continuous external management and funding. The adverse impacts of lethal control, and the shortcomings of existing non-lethal measures create the need for novel non-lethal interventions that have minimal adverse impacts, are within the financial and technical reach of subsistence livestock owners, and are easier and less expensive to apply than lethal control (Shivik 2006). The widespread application of insect pheromones in pest management shows that, once their active ingredients have been identified and formulated, deterrents based on chemical signals are low-tech, low-cost, and quick and easy to apply. The results presented here are the first evidence that mammal chemical signals can be formulated and applied in the same way as those of insects.

## 2. Methods

### 2.1 Study areas

Tests were conducted in northern Botswana and central Zimbabwe (Figure 1);

- Traditional subsistence livestock (locally referred to as cattle posts) around the settlements of Gogomogo, Tsutsubeka and Xhoo (GTX) to the east of Maun (−19.913961° 23.223117°). This area suffers severe impacts from wildlife because it is separated from the NG/30 and NG/32 Wildlife Management Areas only by a veterinary cordon fence that is in serious disrepair.
- Two commercial cattle ranches in the northern Hainaveld (HR1 −20.396460° 23.709497°, HR2 −20.275596° 23.670785°) running cattle and a few sheep and goats, and a commercial cattle ranch in the Motopi area (MR) (−20.163434° 24.037963°). All three ranches lie in the Kalahari Acacia-Baikiaea savanna, and in addition to livestock have low densities of large and medium-sized wild herbivores, spotted and brown hyaenas (*Parahyaena brunnea*), leopards, African wild dogs and smaller predators.
- Shangani Holistic Ranch; (−19.752018, 29.399550) (ZR) runs Nguni cattle with management aimed at restoration from serious bush encroachment caused by prolonged over-grazing There are resident leopards and spotted hyaenas, and smaller predators.

**Fig. 1.**
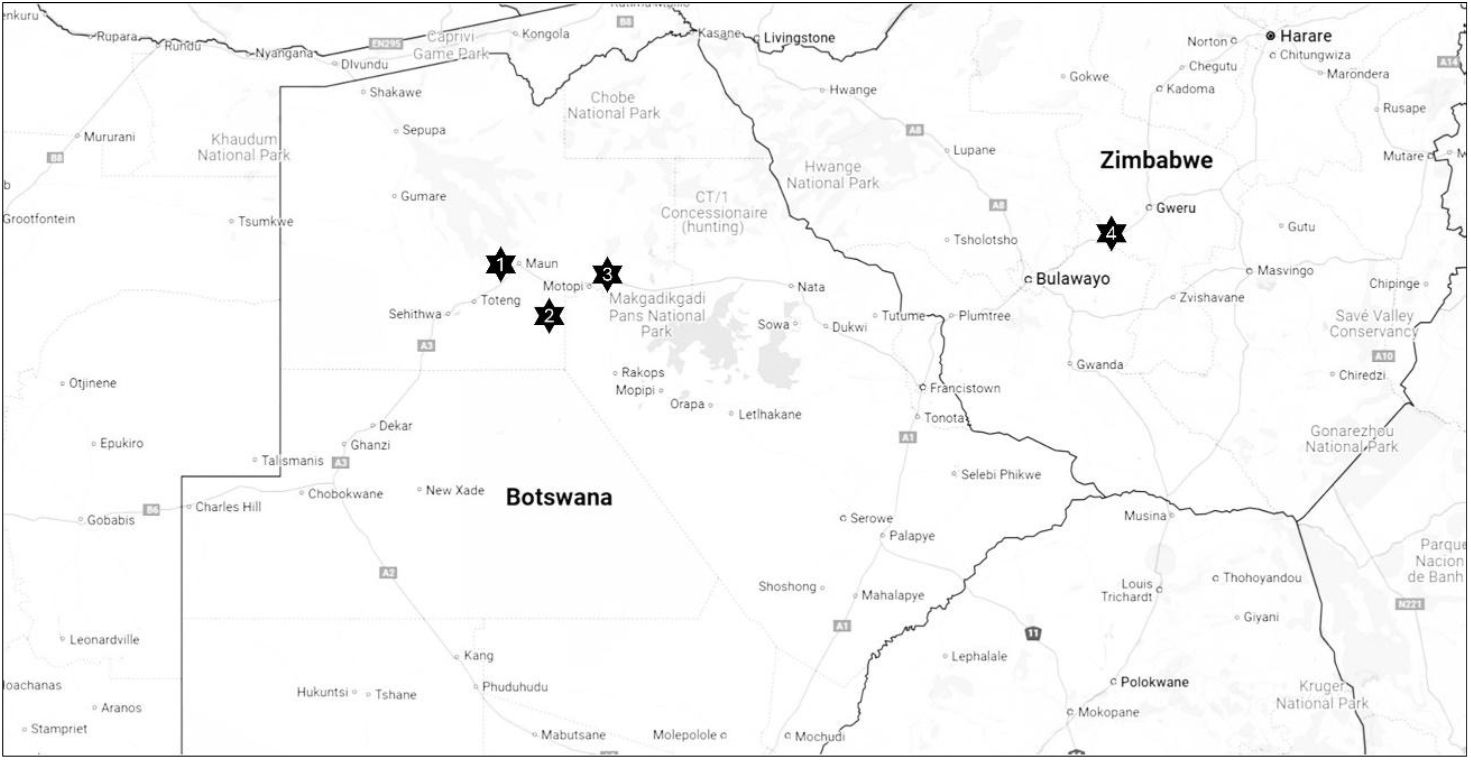
Locations of study sites in Botswana and Zimbabwe. 1 Gogomoga, Tstsubega, Xhoo subsistence cattle posts (GTX), 2 Hainaveld livestock ranches (HR1 and HR2), 3 Motopi cattle ranch (MR), 4 Shangani Ranch.

On the cattle posts and ranches, normal operations continued, including lethal control, trapping and translocation of predators, improvements to fences and enclosures, and creation of new enclosures.

### 2.2 Scent dispensing

3-Mercapto-3-methyl-1-butanol (3M3MB) was obtained from TCI (https://www.tcichemicals.com/JP/en), Carbosynth (https://www.biosynth.com/) and BLDPharm (https://www.bldpharm.com/). Controlled release was achieved by diffusion through silicone septa with the Teflon layer shaved off with a scalpel, on 4 ml or 15 ml open-capped vials (part numbers, Supelco SU860078, Supelco 27115-U, Supelco 27020, Supelco 27088-U respectively) (HR1, MR, ZR), and polyethylene kitchen cling wrap (Glad Cling Wrap, Clorox Africa, Bryanston) on 15 ml open-capped vials (MR, HR2, GTX) (Figure 2). Silicone septa and cling-film on 15 ml vials both generated emission rates of approximately 20 ng/s at 40°C, which is close to the maximum daytime temperature. The odour of 3M3MB was clearly detectable to the human nose 5-10 cm from the vial, but not detectable (to humans) 1 m away.

**Fig. 2.**
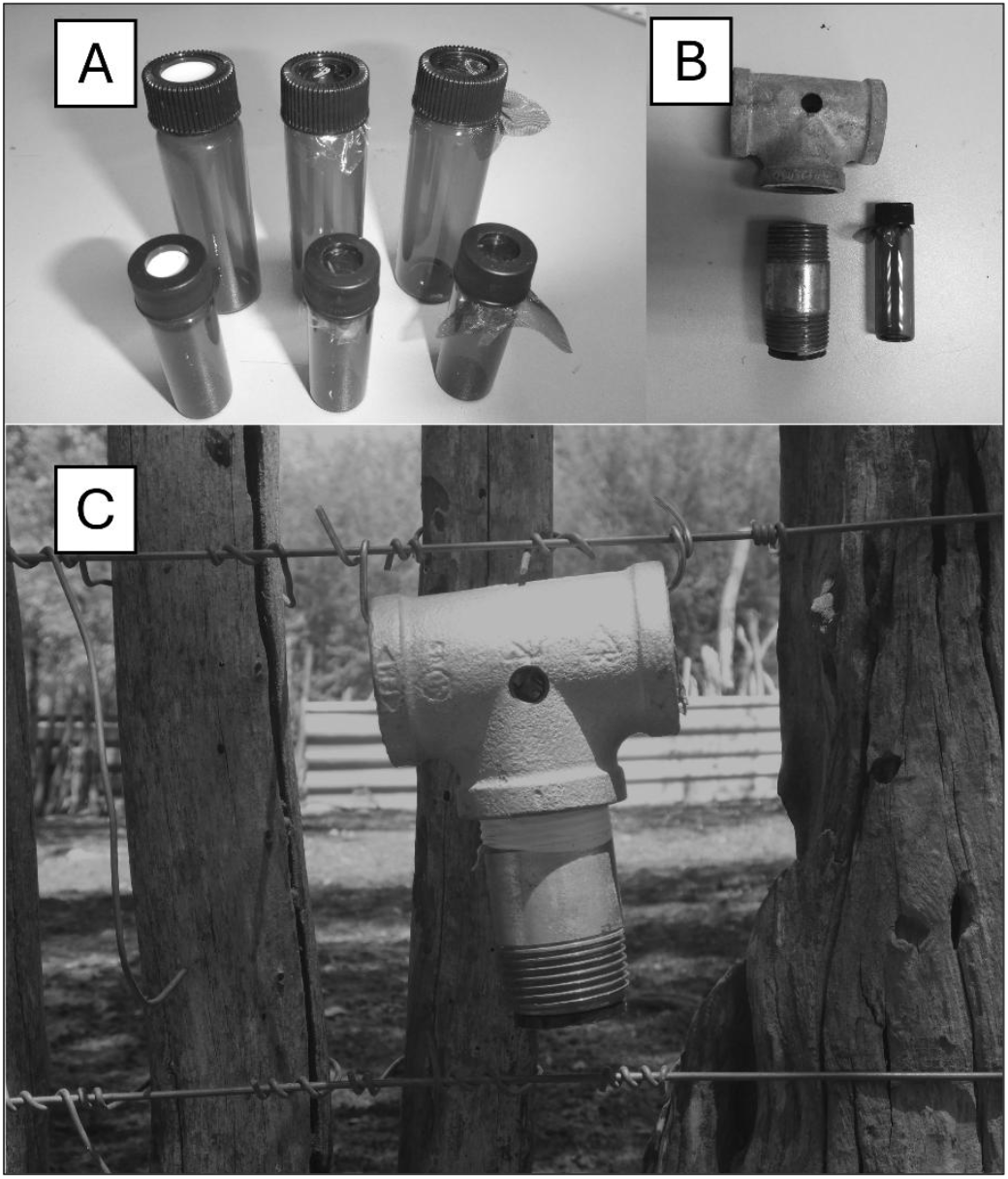
A Controlled release vials; back row 15ml, front row 4ml. Left to right, with; silicone septum, cling-film, cling-film and gauze fabric. B Housing and vial before assembly. C Housing with vial inside wired to a kraal fence.

Increments in release rate were created by increasing membrane area (15 ml *vs* 4ml vials) and increasing the number of dispensers at a site.

To release 3M3MB at kraals, fences and gates, the controlled-release vials were protected by metal housings made from 25 mm galvanised pipe connectors The housings with vials inside were wired to fences about 30 – 50 cm off the ground (Figure 2). For release from animal-borne collars the vials were inserted into holders made from electrical trunking or CPVC water-pipe, that were tied around the animals’ necks with cotton rope (Figure 3).

**Fig. 3.**
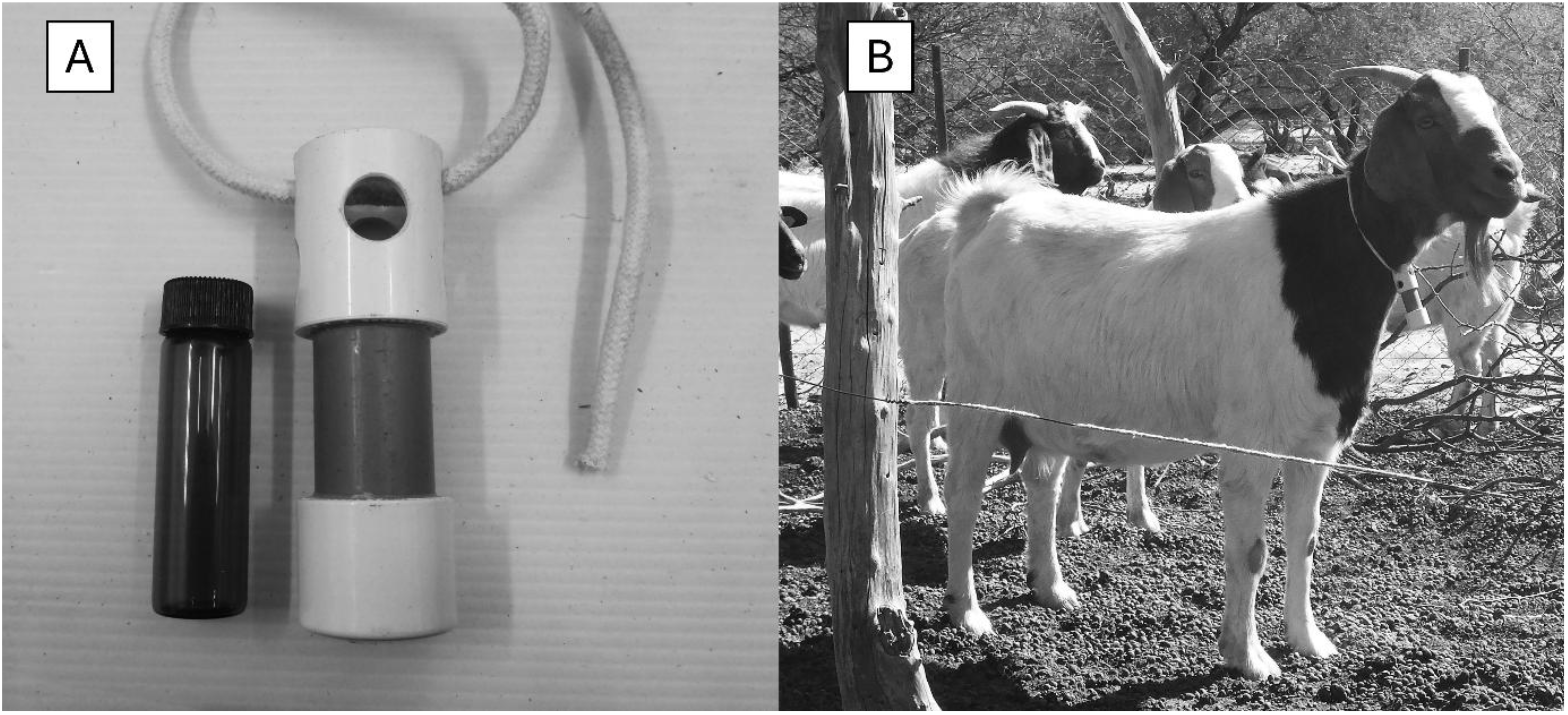
A 15ml dispenser vial and collar for sheep and goats. B A goat wearing a predator deterrent collar.

### 2.3 Camera trapping

The presence of predators and their immediate reactions to 3M3MB were monitored with camera traps. In Botswana we used Reconyx XR6s, Browning Strike Force Pro XDs (Model BTC-5PXD), Browning Patriots (Model BTC-Patriot-FHD) and Browning Spec Ops Elites (model BTC-BE-HP5) (TrailcamPro https://www.trailcampro.com/) mounted 30 – 60 cm off the ground, and set for maximum PIR sensitivity, maximum infrared illuminator power, and 30s videos for Reconyx, 20s videos for Brownings). At Shangani, still images were captured by Cuddeback X-Change™ Color (Model 1279) camera traps attached to the outer fences of the kraal about 1m above the ground, aimed away from the fence.

### 2.4 Livestock losses

Livestock losses were reported by the owners.

### 2.5 Deployments

After a kraal owner in the GTX subsistence area reported having to chase a leopard away from a calf kraal on four consecutive nights, seven silicone-topped dispensers in housings were wired to the kraal fence, and the immediate surroundings of the kraal were monitored with four camera traps aimed outwards from the corners of the kraal.

Dispensers were attached to the GTX veterinary cordon fence where spotted hyaenas used gaps under the wire to cross between the wildlife area and the livestock area. Camera traps were mounted on the fence poles, aimed along the fence towards a gap, or on trees approximately 10m from the fence, aimed at gaps. At the most frequently used gap, release rates were increased at approximately monthly intervals over a period of six months.

At the Hainaveld 1 ranch, predator activity was monitored for four-months without 3M3MB, and for a further 4.5 months with 3M3MB being released from dispensers positioned on both sides of two gateways.

At the Hainaveld 2 ranch, predators passing through a gateway were monitored for one month with two empty dispenser housings on each side of the gate, and then 3M3MB was deployed with incrementally increasing release rates over a period of five months. In June 2024, leopards killed 7 sheep and goats out of a flock of about 30, and six of the survivors were fitted with collars emitting 3M3MB in July 2024 (Figure 3).

At the Motopi cattle ranch, 3M3MB was released from dispensers on kraal fences, initially in response to a female leopard and her grown cub killing four calves in three nights in August 2019. Two days later, eight 3M3MB dispensers with 15ml vials were hung approximately 15 m apart at the corners and mid-points of the kraal fence. The dispensers were moved around as the kraals were rebuilt, and calves were moved, and left in place for varying periods determined by the availability of 3M3MB, vehicle breakdowns, and COVID movement restrictions.

At Shangani, in June 2022, 3M3MB was dispensed from eight points (corners and middles of sides) on the perimeters of two shade-cloth kraals with a history of leopard kills and recent sightings of leopards or their spoor. In October 2022, 15 3M3MB dispensers were deployed about 15m apart around a thorn-branch kraal where a leopard was repeatedly breaching the wall and killing calves.

Except for the subsistence kraal, Motopi, and Shangani, where deployments were reactive, empty dispenser housings were in place during the control periods to reduce neophobic responses to their mere presence.

## 3. RESULTS

At the subsistence cattlepost, no videos of leopards were captured following deployment of 3M3MB, but a leopard was shot nearby 10 days later in an incident that was not connected with livestock predation.

3M3MB deterred spotted hyaenas from crossing the veterinary cordon fence between the GTX subsistence area and the neighbouring wildlife areas. The strength of the deterrent effect increased with increasing release rate (Figures 4, 5). https://www.youtube.com/watch?v=f8tkQkksLsA

**Fig. 4.**
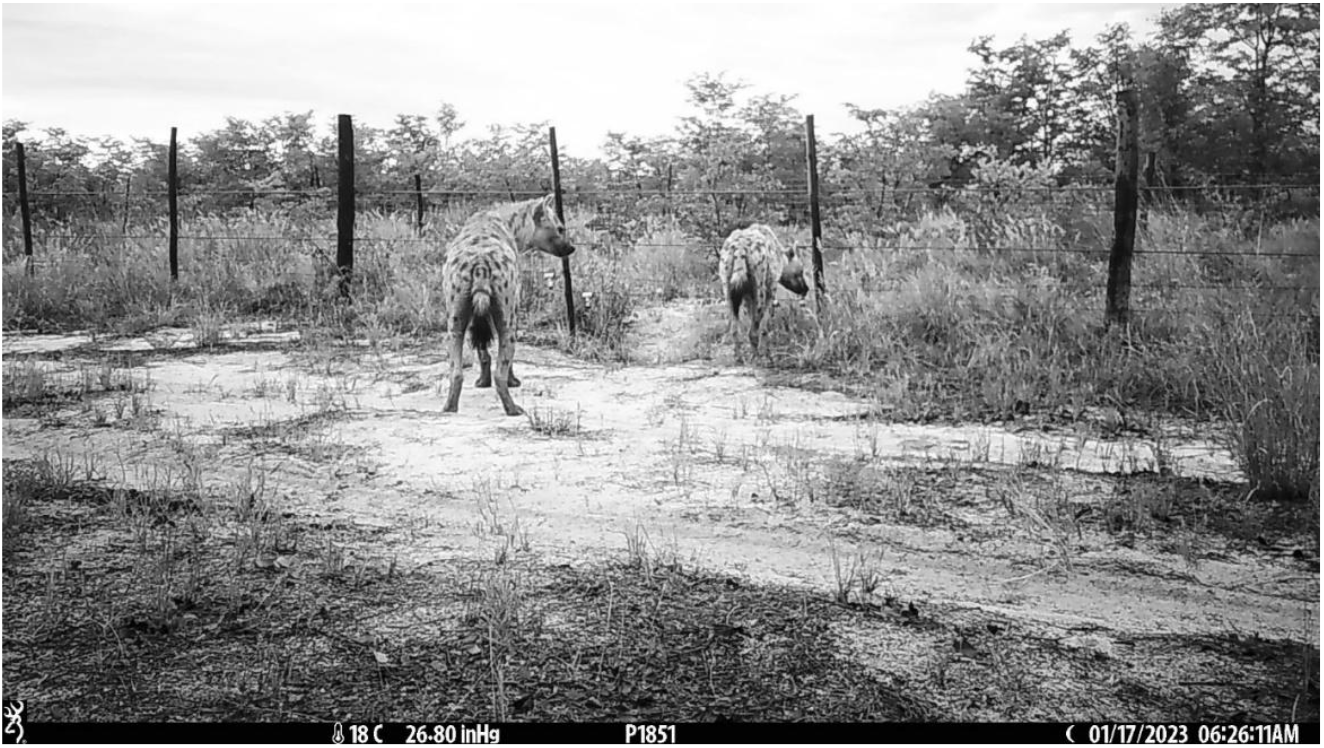
A frame from a video of spotted hyaenas deterred from crossing a veterinary cordon fence by the odour of 3M3MB. The full video is at https://www.youtube.com/watch?v=f8tkQkksLsA

**Fig. 5.**
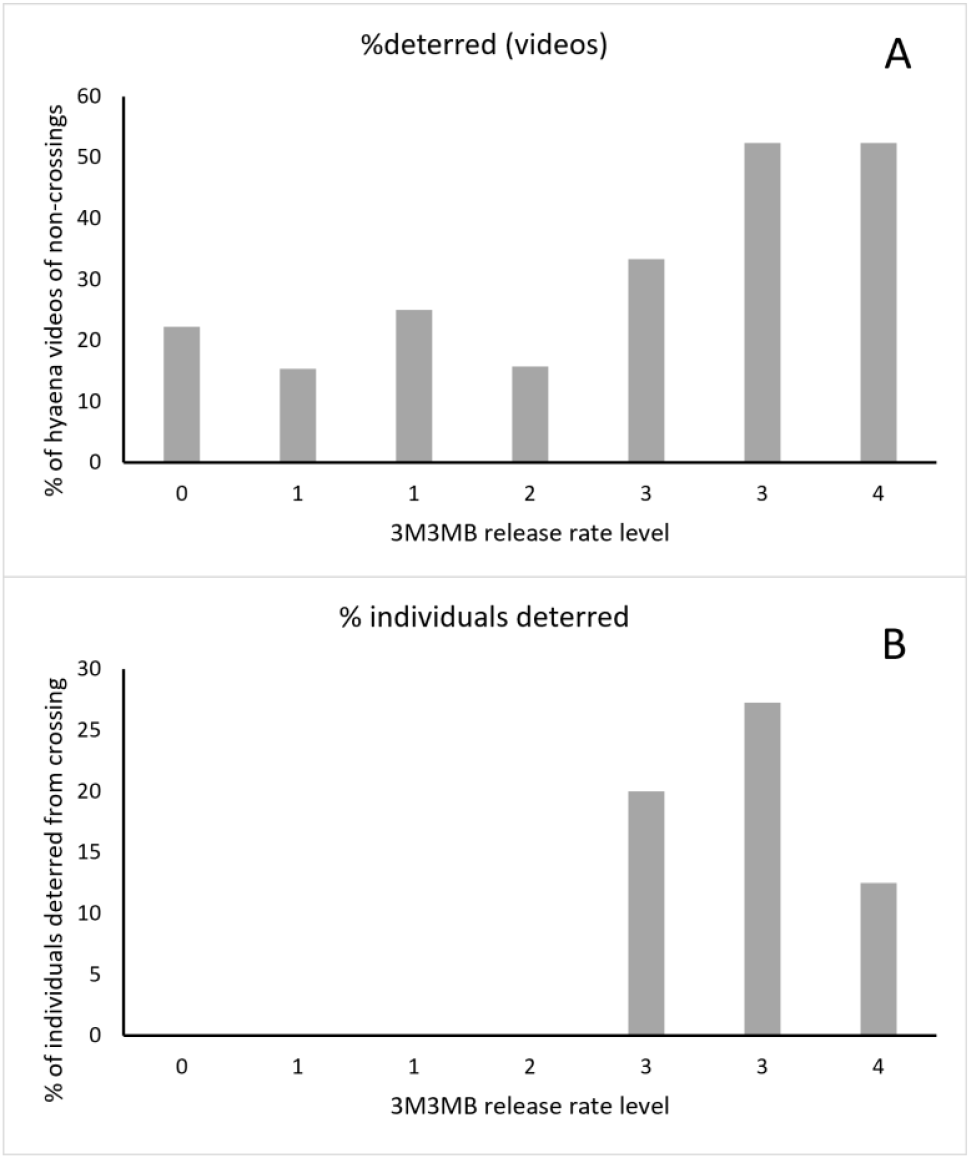
A The percentage of videos of spotted hyaenas at a gap under a veterinary cordon fence that showed hyaenas not crossing the fence, B the percentage of individual hyaenas that did not cross the fence, with increasing release rates of 3M3MB.

On Hainaveld Ranch 1, camera trapping for four-months without 3M3MB yielded seven records of leopards, seven records of spotted hyaenas, and one record of four African wild dogs (*Lycaon pictus*), and a leopard killed a calf but was not recorded on video. Over a further 4.5 months with 3M3MB being released, there were no records of leopards, and only one record each of spotted hyaenas and wild dogs, with no losses of calves.

At Hainaveld Ranch 2, during the one-month control period, three leopards entered through the gate. As the 3M3MB release rate was increased, the number of leopards recorded per month decreased to zero, and then increased again (Figures 6, 7). Reported losses of sheep and goats to leopards dropped from seven in the month before the goats and sheep were given deterrent collars to zero for the next five months.

**Fig. 6.**
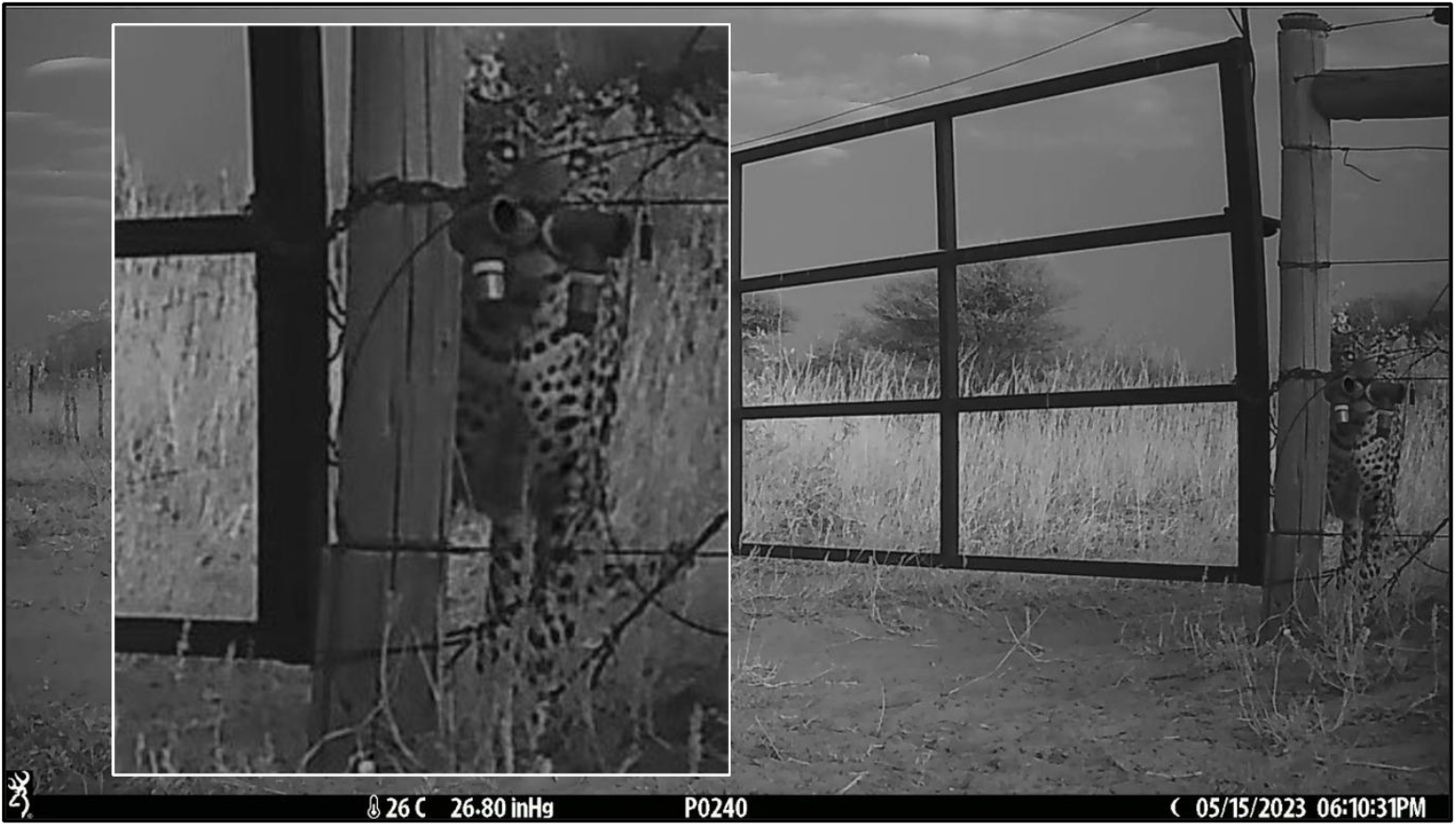
A frame from a video of a leopard sniffing the 3M3MB dispensers before turning back from the gateway. The full video is at https://www.youtube.com/watch?v=PpoTyVJVmDU

**Fig. 7.**
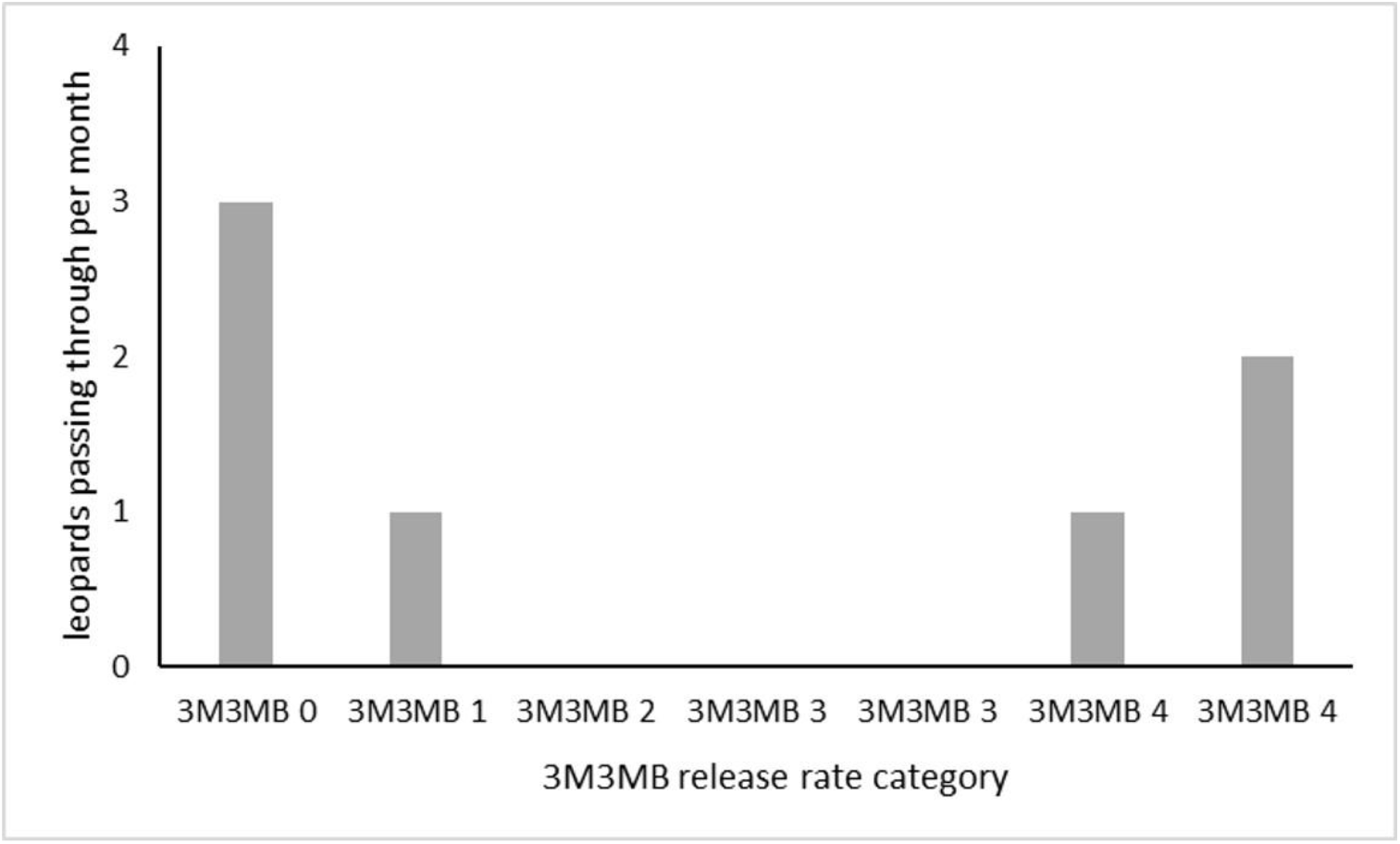
Numbers of leopards passing through a livestock ranch gateway with incrementally increasing rates of 3M3MB release.

At the calf kraals on Motopi cattle ranch, 3M3MB was intermittently released for approximately 18 months over a period of slightly longer than 3 years (Table 1).

**Table 1.**
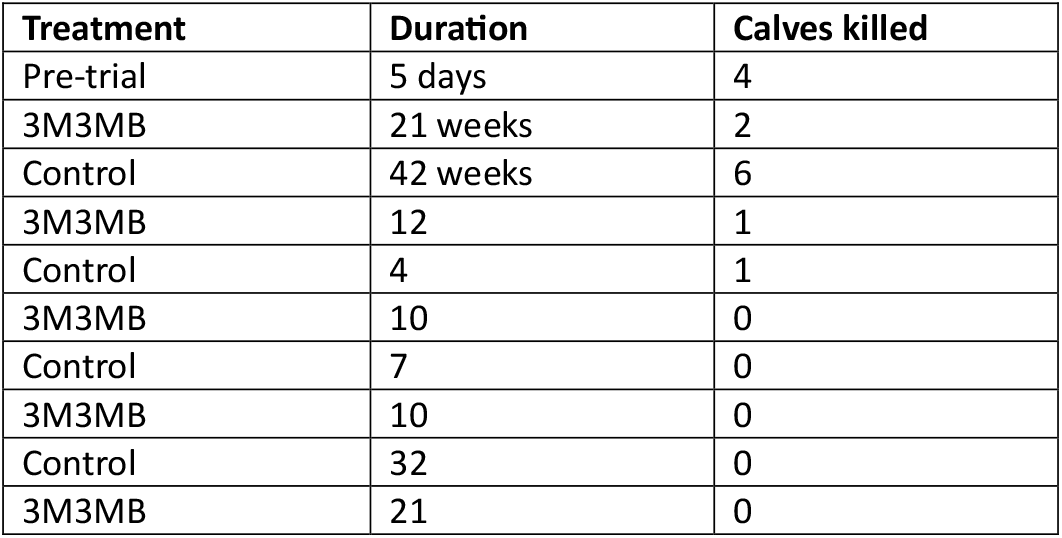
Control and 3M3MB-release periods in chronological order, with numbers of calves taken from a kraal by leopards.

From Nov 2020 the rate of camera-trapped leopard visits declined.to one per month or less, with no livestock losses, irrespective of whether 3M3MB was being released, and so the 3M3MB was removed in August 2022 to re-establish the baseline. In September 2022 the rate of leopard detections at the kraal increased to three per month, and in October 2022 to seven per month, but with no attacks on livestock. In November 2022 the number of leopard records fell again to zero and remained there until February 2023 with no 3M3MB being released. In March 2023, a leopard killed two calves and there was a spike to 14 for the month in camera trap videos of leopards at the kraal (background levels were 0 or 1 per month). Two days after the second calf was killed, eight 3M3MB dispensers were deployed on the fences of the calf kraals, and six extra camera traps were added to the four already at the kraal. At the same time the farmer reinforced the perimeters of the calf kraal with diamond mesh. Over the following 31 days, a leopard was captured on 237 videos of 54 incidents, but despite this high level of activity, no calves were killed. The leopard killed a baby donkey in a kraal with no repellent on 21 April 2023 and was trapped and relocated on 24 April 2023. The repellent and extra cameras were removed on 16 May 2023. On 8 September 2023 a leopard killed two more calves, but killing stopped although no 3M3MB was available. Overall, leopards made 0.2 kills per month when 3M3MB was present and 0.6 kills per month when 3M3MB was absent. The value of the livestock saved was approximately six times the cost of the repellent.

When 3M3MB was present, spoor tracking by Motopi ranch staff showed leopards turning back from the kraal or skirting around it approximately 50 m away. Kraal staff also reported that there were fewer hyaena tracks and less hyaena activity when 3M3MB was present.

At Shangani, in June 2022, camera traps captured images of two leopards just outside shade-cloth kraals where 3M3MB had been deployed, but no livestock were attacked despite the shade cloth fence providing no physical barrier. In September-October 2022, a leopard killed four calves in five weeks at a thorn-branch kraal, and 11 days after the fourth calf was killed 3M3MB and camera traps were deployed for the following month, during which no leopard images were captured, or calves killed. After the 3M3MB was removed, two calves were killed within 10 days.

## 4. Discussion

Camera traps are the best available means of recording the presence of predators and their immediate reactions to deterrents, but the strongest repellent effects will result in animals not being recorded at all because they will not enter the cameras’ detection zones, which extend a maximum of about 20m in front of the camera over a field of view about 45° wide (Apps & McNutt 2018 a, b). Notwithstanding this shortcoming, camera trap videos of predators turning back after sniffing 3M3MB are the most convincing evidence of its effects; neither coincidence nor regression to the mean (see below) can explain animals doing U-turns when they encounter an odour. (Figures 4, 6).

Odour deterrents in collars on goats or sheep provided protection from leopards for sheep and goats (this study), and from caracals for goats (Apps unpubl.); with losses stopping immediately and completely, even though on the Hainaveld 2 ranch at least three leopards continued to be captured by camera traps, and leopards killed uncollared calves within 100 m of the collared sheep and goats. Odour deterrents protected whole flocks, even though only about one third of the animals had a collar. Eye-spots or crosses painted onto cow’s rumps protect the painted animals, but not other herd members (Radford et al. 2021). In Europe, pilot trials are being run on sheep collars that emit wolf “pheromones” (https://gettotext.com/herd-protection-idea-when-the-sheep-smells-like-a-wolf-news/, https://uk.irsea-institute.com/wolf-project/, https://www.iamexpat.ch/lifestyle/lifestyle-news/swiss-sheep-given-scented-collars-help-stop-wolf-attacks, https://www.telegraph.co.uk/world-news/2023/12/01/sheep-collar-pheromones-scares-wolf-predators-switzerland/) but at the time of writing (Dec 2024) no results on their effectiveness have been published.

The repellent effect of 3M3MB on spotted hyaenas was unexpected, because 3M3MB has not been found in spotted hyaena odours (Hofer et a 2001; Apps unpubl.) but it may well be an interspecific cue that alerts the hyaenas to the presence of a potentially dangerous competitor. Interspecific responses to odours are widespread and common, but seriously neglected (Apps et al. 2019).

Any of the quantitative results presented here could have been coincidence; a leopard might have moved away from livestock just before or just after 3M3MB was released for reasons unconnected with the 3M3MB, including control measures by livestock owners such as the use of diamond-mesh fence by the rancher at Motopi. Nonetheless, coincidence becomes less and less convincing the more instances there are, and at some point the evidence from case studies, with its limitations, becomes sufficient to justify investment in rigorously-designed, large-scale trials.

Overall, livestock predation is rare, and when 3M3MB was deployed in reaction to predation, the subsequent declines in livestock losses could have been due to regression to the mean, although the coincidences in timing between 3M3MB deployment and predation stopping suggests otherwise. To evaluate the relevance of regression to the mean would need comparable series of data on losses of livestock and camera trap detections of predators in the absence of deterrents, and these are not currently available.

At both of the sites where the release rate of 3M3MB was systematically increased, the strength of its effects increased with release rate, at one site with leopards and at the other with spotted hyaenas. Dose-response studies are common in work in insect pheromones, and there is some work on lures and baits for mammals (e.g. Jackson et al. 2018) but they are very scarce for mammal semiochemicals, even for laboratory and farm species (Dorries et al. 1995; Leinders-Zufall et al. 2000; He et al. 2010; Li et al. 2013), and these results are the first for a chemical signal from a wild mammal in the wild. In epidemiology, dose-response relationships are evidence of cause and effect (e.g. Shimonovich et al. 2021) but mammal semiochemistry, especially in the wild, is still at the level of Snow and Potts (Bradford Hill 1965).

The reason why the number of leopards passing through the HR2 gate increased at the highest rate of 3M3MB release is unknown. Habituation or another form of learning is plausible, because 3M3MB had been continuously released there for four months. There may also be individual differences in sensitivity to the deterrent, or in motivation, as was apparently the case with spotted hyaenas at the veterinary cordon fence (Figure 5 B); some individuals were deterred from crossing and others were not. While there are a few mammalian chemical signals whose activity depends on concentration, (Review Apps 2013), only two; a putative rat (*Rattus norvegicus*) oestrus signal (Nielsen et al. 2011) and the rabbit (*Oryctologus cuniculus*) mammary pheromone (Coureaud et al. 2004), have been shown to have effects that increase and then decrease with increasing concentration. No peak in response has been demonstrated for the other species where different concentrations have been tested; galagos (*Galago crassicaudatus*) (Crewe et al. 1980), tree shrews (*Tupaia belangeri*) (Von Stralendorff 1982, 1987), and mice (*Mus musculus*) (Ninomiya and Kimura 1990; Li et al. 2013). As far as I know, there is no relevant background information on individual differences in response to odours, or on how long it might take a wild animal to habituate to intermitted exposure to an odour or learn that it is irrelevant.

In domestic cats (*Felis silvestris catus*) 3M3MB’s metabolic precursor is produced as a spill-over from the cholesterol pathway (Miyazaki et al. this vol), which may make 3M3MB an honest signal of a felid’s body condition, and if it is, a higher release rate would signal the presence of a more dangerous competitor to both other leopards and to spotted hyaenas.

Laboratory animals respond to single components of social odours (review Apps 2013) and wild African predators and red foxes (*Vulpes vulpes*) have been shown to scent-mark in response to simplified mixtures or single components of their social odours (Whitten et al. 1980, Apps et al. 2017; Apps & McNutt 2023), but this is the first study that shows wild animals’ movements and hunting being affected by a single signal component, and that it offers protection against livestock predation.

As the success of insect pheromones shows, the field application of deterrents based on chemical signals is low-tech, low-cost, and quick and easy. Formulation, packaging, marketing, and distribution are within the technical reach of small local businesses, so that wildlife conflict amelioration can be embedded into local economies. If predator deterrents reduce livestock losses to predation by an extremely conservative 10% (reduction in losses at the Motopi calf kraal was around 66%, and with collared sheep and goats 100%) then the savings in South Africa will be ZAR100 million in direct costs and ZAR450 million total (US$5.4 million and US$24.3 million respectively) *per annum*. The costs of deterrents would be recovered in months rather than the years required for predator-proof kraals (Weise et al. 2018). Savings in the USA would be US$14 million *per annum*. Because the value of the livestock saved will be higher than the cost of the deterrents, deterrent-based conflict reduction will not need continual financial subsidies, unlike other non-lethal measures. Of all the non-lethal conflict interventions, odour-based deterrents are the only ones with any prospect of being cheaper and easier to apply than bullets, snares and poisons, and the only ones likely to be spontaneously adopted by African farmers and integrated into local economies rather than depending permanently on inputs of foreign money and expertise. This demonstration that predators’ behaviour is affected by a single component of their odours, and that this compound protects livestock in real-life trials, opens the door to the practical application of artificial mammalian chemical signals as minimally invasive, ecologically benign, economically sustainable wildlife management tools.

## Acknowledgements

Funding and donations in kind for the work reported here were provided by; the Leopardess Foundation, WWF Netherland InnoFonds *via* Stichting Spots, the WildIze Foundation, Oppenheimer-Generations Research and Conservation, the Eppley Foundation for Research, Spectral Works, and Ceva Wildlife Research Fund. Dr Marius Viljoen, and Rra Ditso Ntloyathuto, allowed us to work on their land, and the Village Development committees and people of Tsutsubega, Gogomoga and Xhoo provided information on predator activity, and access to their cattle posts. Rra Johane Masene provided technical support in the field in Botswana and the Shangani Ranch Research team assisted in setting up and monitoring the project in Zimbabwe. The work was carried out under research permit EWT 8/36/4 XXXVIII (15) granted by the Ministry of Environment, Wildlife and Tourism and the Office of the President of Botswana.

